# The oomycete MAMP, arachidonic acid, and an *Ascophyllum nodosum*-derived plant biostimulant induce defense metabolome remodeling in tomato

**DOI:** 10.1101/2022.10.11.511777

**Authors:** Domonique C. Lewis, Timo van der Zwan, Andrew Richards, Holly Little, Gitta L. Coaker, Richard M. Bostock

## Abstract

Arachidonic acid (AA) is an oomycete-derived MAMP capable of eliciting robust defense responses and inducing resistance in plants. Similarly, extract (ANE) from the brown seaweed *Ascophylum nodosum*, a plant biostimulant that contains AA, can also prime plants for defense against pathogen challenge. A previous parallel study comparing the transcriptomes of AA and ANE root-treated tomato demonstrated significant overlap in transcriptional profiles, a shared induced resistance phenotype, and changes in accumulation of various defense-related phytohormones. In this work, untargeted metabolomic analysis via liquid chromatography-mass spectrometry was conducted to investigate the local and systemic metabolome-wide remodeling events elicited by AA- and ANE-root treatment in tomato. Our study demonstrated AA and ANE’s capacity to locally and systemically alter the metabolome of tomato with enrichment of chemical classes and accumulation of metabolites associated with defense-related secondary metabolism. AA and ANE root-treated plants showed enrichment of fatty acyl-glycosides and strong modulation of hydroxycinnamic acids and derivatives. Identification of specific metabolites whose accumulation was affected by AA and ANE treatment revealed shared metabolic changes related to ligno-suberin biosynthesis and the synthesis of phenolic compounds. This study highlights the extensive local and systemic metabolic changes in tomato induced by treatment with a fatty acid MAMP and a seaweed-derived plant biostimulant with implications for induced resistance and crop improvement.

## Introduction

Plants have pre-formed and inducible structural and biochemical mechanisms to prevent or arrest pathogen ingress and colonization. These defenses include barriers such as papillae and ligno-suberized layers to fortify cell walls, and low-molecular weight inhibitory chemicals (Freeman and Beattie 2008). Plants undergo transcriptional changes upon perception of microbe associated molecular patterns (MAMPs) or effectors to induce local and systemic resistance.

The oomycete MAMPs, arachidonic acid (AA) and eicosapentaenoic acid (EPA), are potent elicitors of defense. These eicosapolyenoic acids were first identified as active components in *Phytophthora infestans* spore and mycelial extracts capable of eliciting a hypersensitive-like response, phytoalexin accumulation, lignin deposition, and protection against subsequent infection in potato tuber discs (Bostock et al. 1981; Bostock et al. 1982). Further work demonstrated root treatment with AA protects tomato and pepper seedlings from root and crown rot caused by *Phytophthora capsici*, with associated lignification at sites of attempted infection (Dye and Bostock 2021). AA has been shown to induce resistance, elicit production of reactive oxygen species, and trigger programmed cell death in members of the Solanaceae and other families (Araceli et al. 2007; Bostock et al. 1981; Cook et al. 2018; Dedyukhina et al. 2014; Dye et al. 2020; Knight et al. 2001).

Phaeophyta and Rhodophyta members (red and brown macroalgae) contain numerous bioactive chemicals that can elicit defense responses in plants (Klarzynski et al. 2003; Sangha et al. 2010; Vera et al. 2011). The brown alga, *Ascophyllum nodosum*, is a rich source of polyunsaturated fatty acids, including AA and EPA, which comprise nearly 25% of its total fatty acid composition (Lorenzo et al. 2017; van Ginneken et al. 2011). *A. nodosum* and oomycetes belong to the major eukaryotic lineage, the Stramenopila, and share other biochemical features (e.g., both are rich in β-1,3-glucans). Commercial extracts of *A. nodosum*, used in organic and conventional agriculture as plant biostimulants, may also help plants cope with biotic and abiotic stresses. A proprietary *A. nodosum* extract, Acadian (hereafter ANE; APH-1011-00, Acadian Seaplants, Ltd., Nova Scotia, Canada), has been shown to provide protection against bacterial and fungal pathogens (Ali et al. 2016a). Studies in *A. thaliana* showed ANE induced systemic resistance to *Pseudomonas syringae* pv. *tomato* and *Sclerotinia sclerotiorum* (Subramanian et al. 2011). Investigation into ANE-induced resistance in *A. thaliana* and tomato suggest the role of ROS production, jasmonic acid signaling, and upregulation of defense-related genes and metabolites (Ali et al. 2016b; Cook et al. 2018; Jayaraj et al. 2008; Subramanian et al. 2011). As a predominant polyunsaturated fatty acid in ANE, AA may contribute to ANE’s biological activity.

In a parallel study we demonstrated AA’s ability to systemically induce resistance and ANE’s capacity to locally and systemically induce resistance in tomato to different pathogens (Lewis et al. 2022). Further, we showed that AA and ANE altered the phytohormone profile of tomato by modulating the accumulation of defense-related phytohormones (Lewis et al. 2022). Through RNA sequencing, this same study revealed a striking level of transcriptional overlap in the gene expression profiles of AA- or ANE-root-treated tomato across tested timepoints (Lewis et al. 2022). Gene ontology functional analysis of transcriptomic data revealed AA and ANE enriched similar categories of genes with nearly perfect overlap also observed in categories of under-represented genes. Both AA and ANE treatment protected seedings from challenge with pathogens with different parasitic strategies while eliciting expression of genes involved in immunity and secondary metabolism. The shared induced resistance phenotype and extensive transcriptional overlap of AA and ANE treatments suggested similar metabolic changes may be occurring in treated plants. In the current study, untargeted metabolomic analyses were conducted to assess global effects of root treatment with AA and the AA-containing complex extract, ANE, on the metabolome of tomato plants.

## Materials and Methods

### Plant growth and root treatment

#### Plant materials and hydroponic growth system

Seeds of tomato (*Solanum lycopersicum cv*. ‘New Yorker’) were surface-sterilized and germinated for 10 days on germination paper. Seedlings were transferred to a hydroponic growth system in 0.5X aerated Hoagland’s solution contained in darkened plastic containers and maintained in a growth chamber with the following conditions: light intensity 150 μmol m^-2^ s^-1^, 16-hour photoperiod, 24°C day, 22°C night, 65% relative humidity (Dye et al. 2020). The seedlings are incubated in the growth chamber for approximately 10 days until emergence of two true fully expanded leaves. Seed was obtained from a commercial source (Totally Tomatoes, Randolph, WI).

#### Root treatment and tissue harvest

Fatty acid sodium salts (Na-FA; Nu-Chek Prep, Elysian, MN) were prepared and stored as previously described (Dye et al. 2020). AA stock solution was prepared by dissolving 100 mg of fatty acid salt in 1 mL of 75% ethanol. AA stock solution was subsequently stored in a glass vial at -20°C flushed with N_2_ to minimize oxidation. A proprietary formulation of *A. nodosum* extract (ANE; APH-1011-00; Acadian Seaplants, Ltd., Nova Scotia, Canada) was diluted with deionized water (diH2O) to a 10% working concentration, which was used to prepare treatment dilutions. All chemicals were diluted to their treatment concentrations with sterile diH_2_O. Hydroponically reared, 3-week-old tomato seedlings with two fully expanded true leaves were transferred to 1 L darkened treatment containers with their respective root treatment solutions. Following 24, 48, 72, and 96 hours of root treatment, tomato seedlings were removed from treatment containers, and leaves and roots were excised from shoots and flash frozen in liquid nitrogen. Each sample was the pool of roots or leaves of two seedlings with four replications per tissue, treatment, and timepoint.

### Metabolite extraction

Samples were transported on dry ice and stored at −70 °C until metabolite extraction. Tissue samples were ground in liquid nitrogen using a mortar and pestle and 100 mg was weighed and transferred to a 2-ml bead-beating tube containing four 2.8-mm ceramic beads. All tools and consumables were pre-chilled in liquid nitrogen. After weighing, each sample was removed from liquid nitrogen and kept at −20 °C until addition of extraction solution.

One ml of extraction solution (80 % v/v methanol and 0.1 % v/v formic acid in ultrapure water) was added to each sample which was then vortexed, followed by bead-beating in a bead mill (Bead Mill 24, Fisherbrand) at a speed of 2.9 m/s for one 3-min cycle. After bead-beating, samples were centrifuged at 12k × *g* for 10 min at 4 °C (Accu Spin Micro 17R, Thermo Scientific). Samples were diluted 5-fold using extraction solution and filtered into LC-MS-grade HPLC vials using 0.22-μm PTFE syringe filters. HPLC vials were kept at 4 °C until LC-MS analysis. A blank was prepared by adding 1 ml extraction solution to a bead-beating tube containing beads that was processed equivalently to the samples. In addition, a quality control sample was prepared by combining 20 μl of each of the extracted samples and processed equivalently.

### Liquid chromatography-mass spectrometry (LC-MS) run conditions

Samples were analyzed via high performance liquid chromatography (Agilent 1260 Infinity) and electrospray ionization quadrupole time-of-flight mass spectrometry (Agilent 6530 Q-TOF) controlled by MassHunter software in centroid data mode. Mobile phase A was ultrapure water with 0.05 % (v/v) formic acid and mobile phase B was acetonitrile with 0.05 % (v/v) formic acid. Before starting the run, the column (Poroshell 120 EC-C18; 3.0 mm internal diameter, 50 mm length, 2.7 μm particle size; Agilent), equipped with a guard column (EC-C18; 3.0 mm internal diameter, 5 mm length, 2.7 μm particle size; Agilent), was conditioned for 20 minutes with 95 % mobile phase A and 5 % B. Column temperature was maintained at 40 °C. The sample injection order was randomized, with individual samples being run consecutively in positive and negative mode. The quality control sample was injected at the beginning and end of the run, as well as after every 12 samples throughout the run to check signal and elution stability. Source parameters were as follows: drying gas temperature of 325 °C (positive) and 350 °C (negative), drying gas flow 12 l/min, nebulizer pressure 35 psi, sheath gas temp 375 °C (positive) and 400 °C (negative), sheath gas flow 11 l/min, capillary voltage 3500 V (positive) and 3000 V (negative), nozzle voltage 0 V (positive) and 1500 V (negative), fragmentor 125 V, skimmer 65 V, and octopole 750 V. Acquisition was performed over a mass range of 50 to 1700 m/z using the all-ions MS/MS technique, cycling three different collision energies (0, 10, 30 eV) at an acquisition rate of 3 spectra/s. Simultaneous infusion of a solution of purine and hexakis(1H, 1H, 3H-tetrafluoropropoxy)phosphazine using the reference nebulizer was used throughout the runs for mass calibration. Samples were introduced using 2 μl injections at a flow rate of 0.5 ml/min. A needle wash of 1:1 acetonitrile:water was used between each injection. The mobile phase gradient was: 0 min, 95 % A, 5 % B; 1 min, 95 % A, 5 % B; 10 min, 50 % A, 50 % B; 15 min, 0 % A, 100 % B; 17 min, 0 % A, 100 % B; 17.1 min, 95 % A, 5 % B; 19.1 min, 95 % A, 5 % B.

### Data analysis pipeline

#### Data alignment, deconvolution, and normalization

Positive and negative mode raw data files from MassHunter were analysed separately in MS-DIAL (v. 4.80) before downstream analysis. Tolerances for MS1 and MS2 were set to 0.025 and 0.075 Da respectively (Tsugawa et al. 2015). A representative quality control sample run was used as the reference file to align peaks. For peak detection, the mass slice width was set to 0.1 DA and the minimum peak height was set to 15,000 which was approximately 3 times the noise level observed in the total ion chromatogram. A linear weighted moving average method was used for peak smoothing, with a smoothing level of 3 scans and a minimum peak width of 5 scans. Deconvolution was performed with a sigma window value of 0.5 and an MS/MS abundance cutoff of 10. The adducts permitted were [M+H]+, [M+NH_4_]+, [M+Na]+, [M+K]+, [M+H−H_2_O]+, and [2M+H]+ in positive mode, and [M−H]−, [M−H_2_O−H]−, [M+Cl]−, [M+Na−2H]−, and [M+K−2H]− in negative mode.

#### Quality control, feature annotation, and merging

MS-DIAL data was cleaned in MS-CleanR (*mscleanr*, v. 1.0) in RStudio (v. 1.4.1106) using the following parameters: minimum blank ratio of 0.8, maximum relative standard deviation of 30, minimum relative mass defect (RMD) of 50, maximum RMD of 3000, maximum mass difference of 0.05 and maximum retention time difference of 0.15. MS-DIAL features were clustered by applying a Pearson correlation, with a minimum correlation of 0.8 and maximum p value of 0.05, retaining two features per cluster according to most intense and the most connected peak filters (Fraisier-Vannier et al. 2020). Selected peaks were imported into MS-FINDER (v. 3.52) for annotation (Tsugawa et al. 2016). Mass tolerance for MS1 and MS2 were set to 5 and 15 ppm respectively, the relative abundance cut off set at 1% and the formula finder was configured to use C, H, O, N, P, and S atoms. FooDB, PlantCyc, ChEBI, NPA, NANPDB, COCONUT, KNApSAcK, PubChem, and UNPD were used as local databases. During the final merge step in MS-CleanR, the best annotation for each peak was based on MS-FINDER scores. The normalized annotated peaks list produced by MS-CleanR was used for the final statistical analyses in R.

#### Statistical analysis

All statistical analyses were completed in R (v. 4.0.5) via RStudio (v. 1.4.1106). Statistical analyses were run separately for each tissue type using the same parameters.

#### Multivariate analyses

A partial least squares (PLS) supervised model of the complete log-transformed and Pareto-scaled dataset was done using the *ropls* package (version 1.22.0), with the three treatment groups as the response variables. Ellipses were drawn around treatments using *stat_ellipse* (*ggplot2*) based on a 95% confidence level. This distance type considers the correlation between variables and the ellipses are created around the centroid data point. Heatmaps were created using the log-transformed data within the *ComplexHeatmap* package in R (v. 2.13.1), with hierarchical clustering according to the complete-linkage method and Euclidean distance measure across columns and rows, displayed as dendrograms.

#### Pairwise multivariate analyses

Pairwise multivariate analysis was performed across all time points between ANE and H_2_O, and between AA and H_2_O, using an orthogonal projections to latent structures discriminant analysis (OPLS-DA). OPLS-DA models were generated using the *ropls* package, with the predictive components set to 1 and orthogonal components to 7. S-plots were generated following sample sum normalization and Pareto scaling via calculation of p1 (covariance) and pcorr1 (correlation) of the OPLS-DA scores using the *muma* package (v. 1.4) source code within R. Chemical class enrichment analysis was achieved using ChemRICH (version 1.0 August 2020) for each two-treatment comparison at each time point within R using the source code (Barupal and Fiehn 2017). A student’s t-test of the signal was conducted to generate p values and effect size. The effect size in each case represents the difference compared to the water control treatment group.

## Results

### Local and systemic metabolomic analysis of AA- and ANE-induced plants

Previous transcriptomic work revealed a high level of congruency in differentially expressed genes in AA- and ANE-treated tomato seedlings compared to H_2_O-treated controls (Lewis et al. 2022). To further this line of investigation, the metabolomic profiles of AA- and ANE-root-treated plants were compared to H_2_O after 24-, 48-, 72-, and 96-hours exposure to their respective treatments. Locally-treated roots and distal leaves were harvested, flash frozen, extracted for metabolites, and subsequently analyzed via LC-MS, which with underivatized samples primarily captures the nonvolatile metabolome (Tohge and Fernie 2015) (**Fig. 1A**).

**Figure 1.**
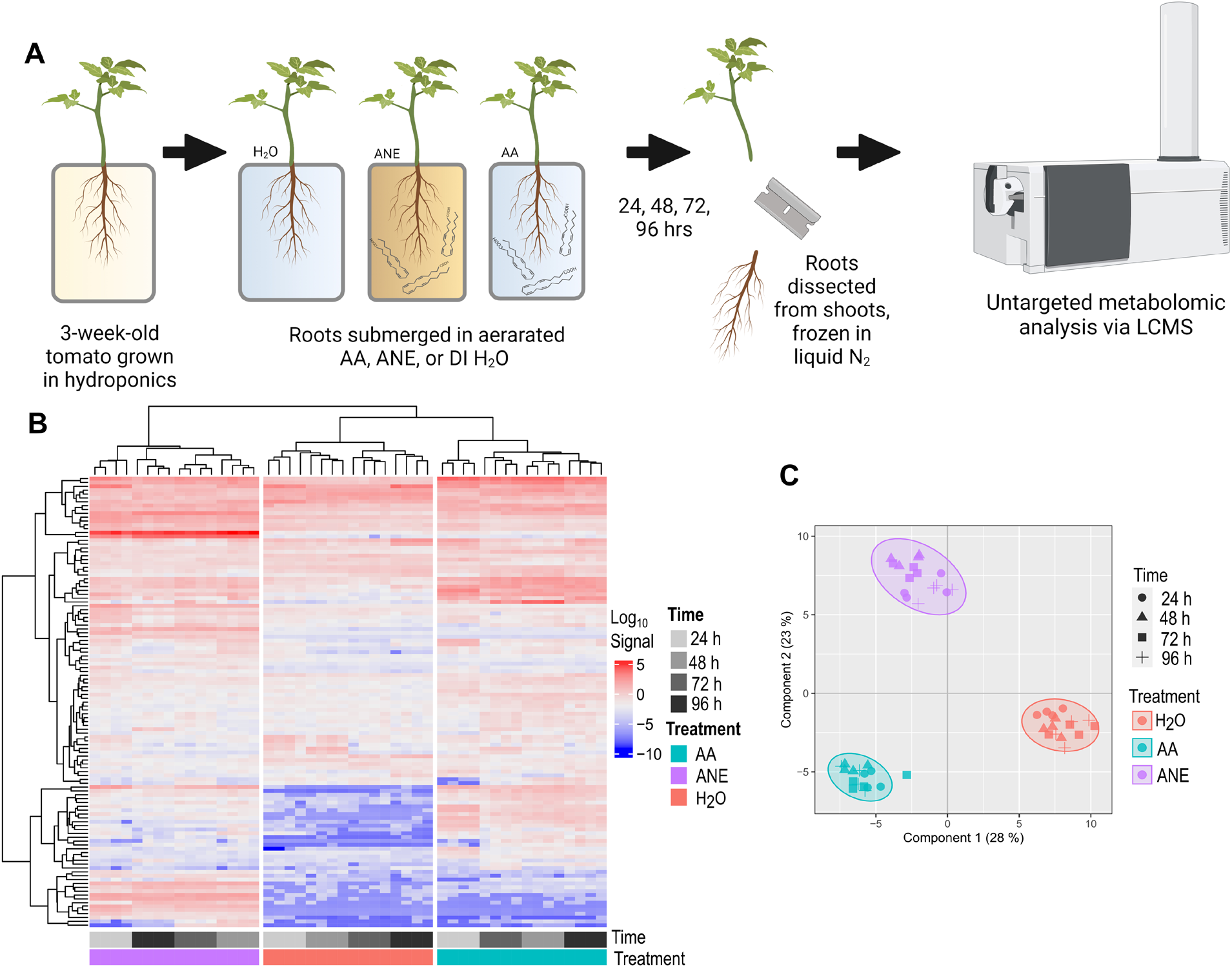
**(A)** Experimental procedure for the untargeted metabolomics study. Tomato roots were treated with 10 μM AA, 0.4% ANE, or H_2_O. Following 24, 48, 72, and 96 hours of root exposure to their respective treatments, plants were harvested, root and leaf tissue dissected from shoots, and the collected tissue flash frozen in liquid nitrogen. Harvested tissue was then subjected to metabolite extraction and liquid chromatography/mass spectrometry. **(B)** Log10 signal of metabolites from root tissue of 10 μM AA-, 0.4% ANE-, or H_2_O-treated plants sampled at 24, 48, 72, and 96 hours or treatment. Features visualized in the heatmap are filtered from the total dataset having an adjusted p-value < 10^−6^ and a fold change > 5. Heatmap data is log10 transformed and hierarchical clustering was conducted across rows and columns. **(C)** Partial least squares score plots of tomato root metabolites after 24, 48, 72, and 96 hours of treatment with 10 μM AA, 0.4% ANE, or H_2_O. Ellipses indicate the 95% confidence intervals. Score plots show variance of 4 biological replicates performed per timepoint and treatment.

Partial least squares (PLS) score plots of tomato root tissue revealed distinct clustering by treatment irrespective of timepoint (**Fig. 1C**). No overlap was observed in the 95% confidence ellipses for any treatment group. Likewise, heatmap visualization of the log_10_ signal of metabolites showed clear clustering of metabolomic profiles by treatment (**Fig. 1B**). Features displayed in the heatmap were filtered from the total dataset with a p-value < 10^−6^ and absolute fold change > 5 in roots.

Less defined clustering was observed in PLS score plots of distal leaf tissue across sampled timepoints (**Fig. 2B**). Ellipses representing the 95% confidence interval for both H_2_O and AA treatments both partially overlap with the ANE treatment group. Similarly, a heatmap depicting metabolite log_10_ signal showed more diffuse clustering by treatment (**Fig. 2A**). Visualized metabolites from leaves displayed in the heatmap used a p-value < 0.001 and an absolute fold-change > 2. These findings are reflective of distal tissue (leaves) not directly treated with either elicitor. Changes in the distal leaves were not as robust as in the directly treated roots, likely due to diminution of systemic signals that effect metabolic changes throughout the plant.

**Figure 2.**
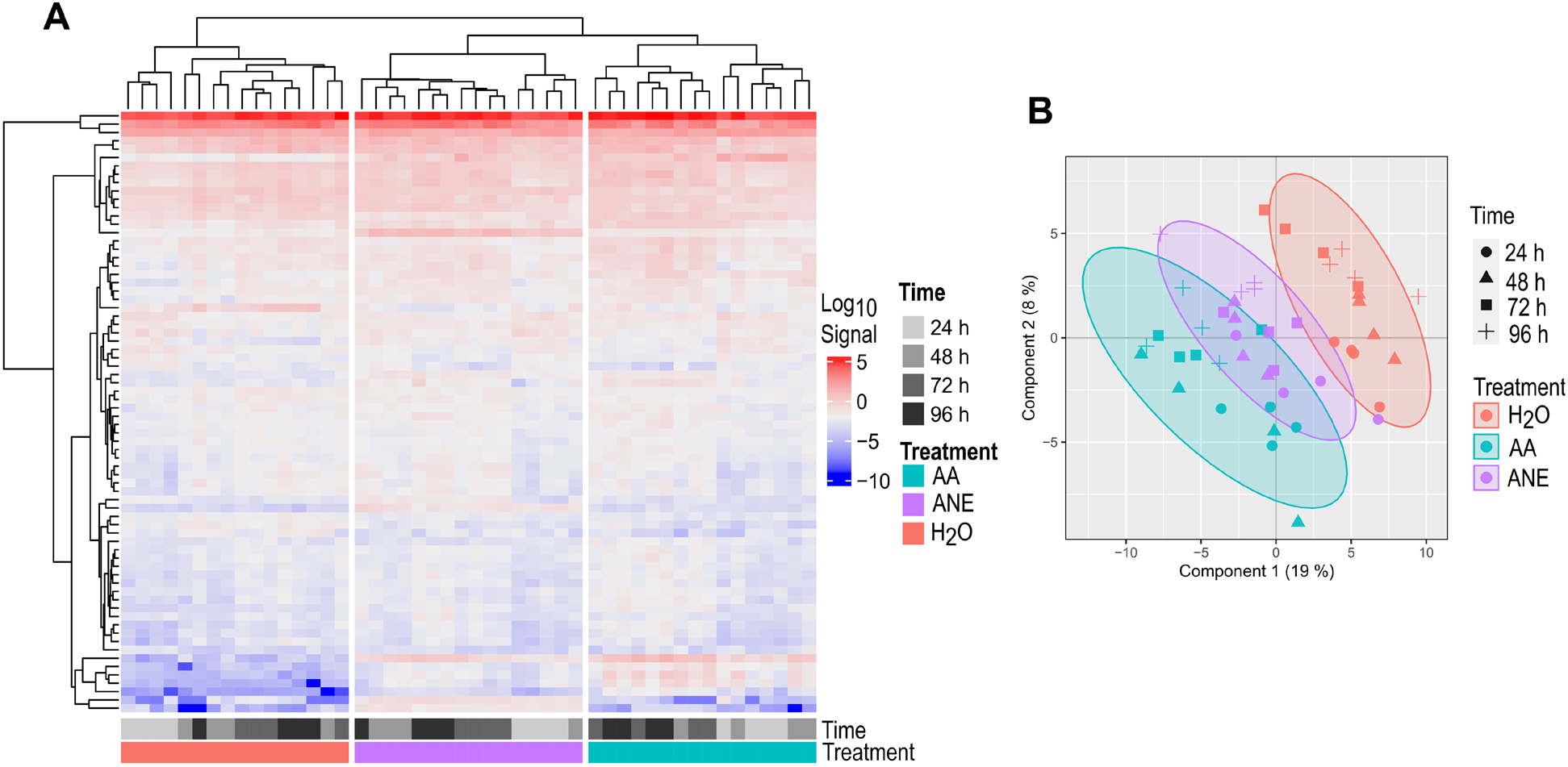
**(A)** Log10 signal of metabolites from leaf tissue of 10 μM AA-, 0.4% ANE-, or H_2_O-treated plants at 24, 48, 72, and 96 hours of treatment. Features visualized in the heatmap are filtered from the total dataset having an adjusted p-value < 0.001 and a fold change > 2. Heatmap data is log10 transformed and heirachical clustering was conducted across rows and columns. **(B)** Partial least squares score plots of tomato leaf metabolites after 24,48, 72, and 96 hours of treatment with 10 μM AA, 0.4% ANE, or H_2_O. Score plots show variance of 4 biological replicates performed per timepoint and treatment. Ellipses indicate the 95% confidence intervals.

An assessment of the total annotated features across the metabolomic analysis revealed shared and unique annotated features between roots and leaves (**Supplementary Fig. S1A**). Roots and leaves share 44 features with leaves displaying the largest number of unique metabolic features (**Supplementary Fig. S1A**). There were 330 unique identified features in leaves compared to 223 features unique to root tissue (**Supplementary Fig. S1A**). A comparison of differential metabolic features for AA- and ANE-treated plants compared to the H_2_O control revealed robust overlap for both roots and leaves (**Supplementary Fig. S1B and S1C**). AA- and ANE-treated roots shared 68 differential metabolic features, with AA and ANE treatments possessing 37 and 29 differential features unique to each, respectively (**Supplementary Fig. S1B**). Less overlap was observed in leaves with 39 shared differential metabolites, with AA- and ANE-root treated plants displaying 34 and 19 uniquely differential metabolites, respectively (**Supplementary Fig. S1C**).

### Chemical enrichment analysis of AA- and ANE-induced plants

Chemical enrichment analyses were conducted to identify classes of metabolites whose accumulation was locally or systemically altered in AA- and ANE-root-treated tomato seedlings. Enrichment analyses of metabolites whose mean signal was significantly changed in AA- or ANE-treated plants compared to H_2_O identified numerous affected chemical classes **(Fig. 3 and Supplementary Fig. S2)**. These changes were most robust in directly treated roots compared to distal leaves. Treatment of tomato seedlings with AA showed strong modulation of metabolomic features classified as triterpenoids and linoleic acid and derivatives in roots. AA-treated roots also showed increases in hydroxycinnamic acids and derivatives and fatty acyl glycosides of mono- and disaccharides. ANE-treated roots showed modulation in the accumulation of triterpenoids, steroidal glycosides, and hydroxycinnamic acids and derivatives. Similar to AA-treated plants, the roots of plants treated with ANE also showed increases in metabolites classified as fatty acyl glycosides of mono- and di-saccharides.

**Figure 3.**
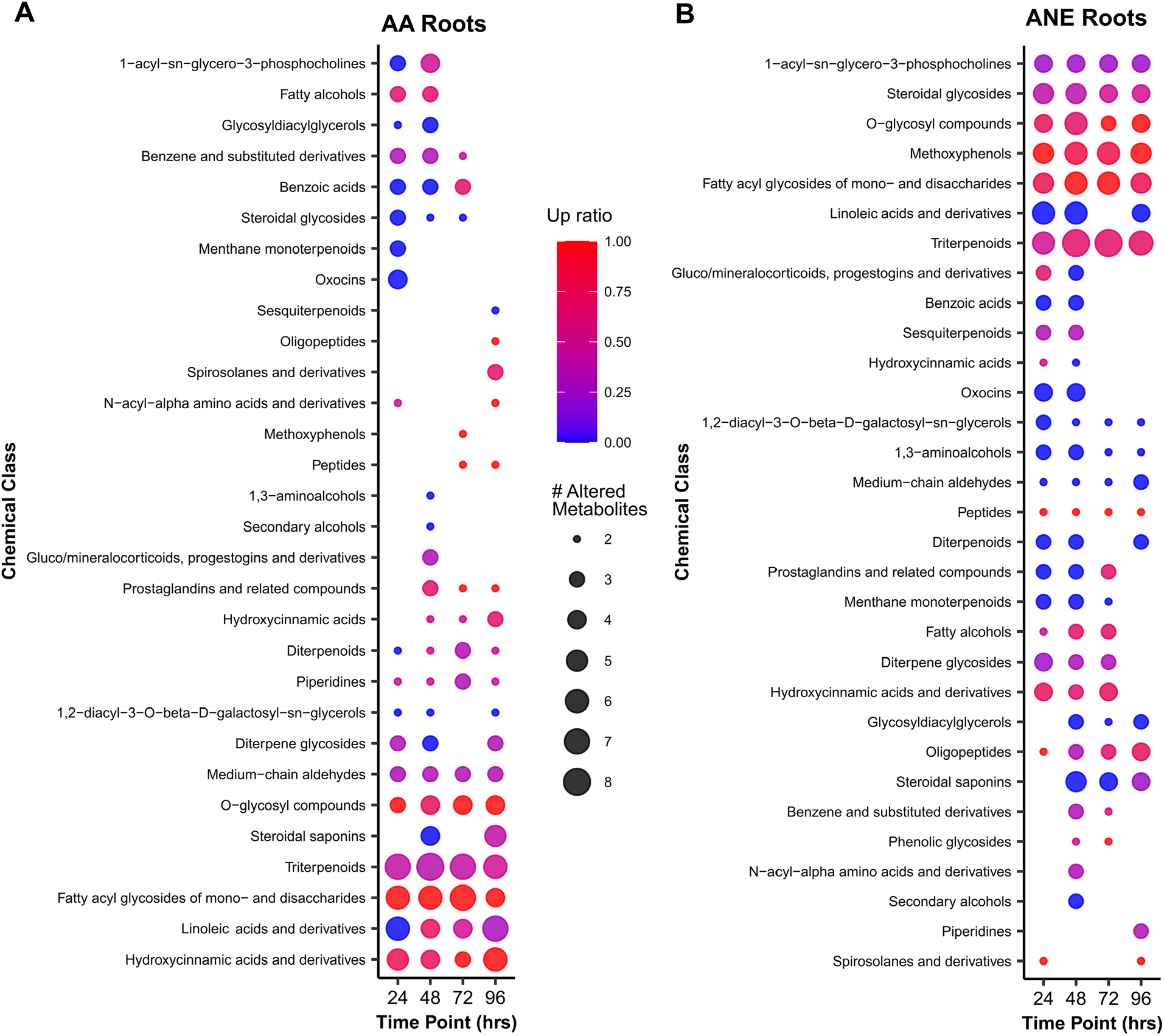
Dot plot illustrating chemical enrichment analysis of significantly altered metabolite clusters in roots of **(A)** AA- and **(B)** ANE-root-treated plants compared to H_2_O after 24, 48, 72, 96 hours of treatment. Dot sizes indicate the number of altered metabolites per identified cluster. Dot color scale indicates the ratio of up (red) and down (blue) compounds in AA- and ANE-treated plants compared to H_2_O. Up ratio represents the proportion of increased/decreased metabolites compared to H_2_O.

Although less striking than the chemical enrichment analysis of roots, leaf tissue of root-treated plants did reveal an altered metabolome **(Supplementary Fig. S2)**. These changes in metabolite accumulation occurred most prominently at 96 hours, the latest tested timepoint.

Increases in sesquiterpenoids and steroidal saponins were seen in leaves of AA-treated plants at 96 hours. A mix of accumulation and suppression of terpenoids and an increase of methoxyphenols was observed in the leaves of ANE-root-treated plants.

### Specific metabolomic features modulated by AA- and ANE-root treatments

Chemical enrichment analyses broadly revealed classes of metabolites that were induced or suppressed in AA- or ANE-treated plants. To examine which specific variables (metabolites) provide the strongest discriminatory power between the two treatment groups, a two-group comparative supervised multivariate analysis, orthogonal projections to latent structures discriminant analysis (OPLS-DA), was utilized. OPLS-DA score plots show strong between-group variability discrimination between AA and ANE treatment groups compared to the H_2_O control across all tested timepoints with the x-axis describing the inter-treatment variability, and the y-axis showing the intra-treatment variability **(Supplementary Fig. S3)**. S-plots derived from OPLS-DA were examined for both AA and ANE treatments in pairwise comparison with H_2_O control. S-plots of OPLS-DA revealed that treatment with AA or ANE induced shared changes in the levels of several defense-related metabolites in roots **(Fig. 4)**. Variables (metabolites) with the most negative and positive correlation and covariance values are the most influential in the model. These metabolites are located on either tail of the S-plot (upper-right and lower-left) and contribute most greatly to the separation between treatment groups

**Figure 4.**
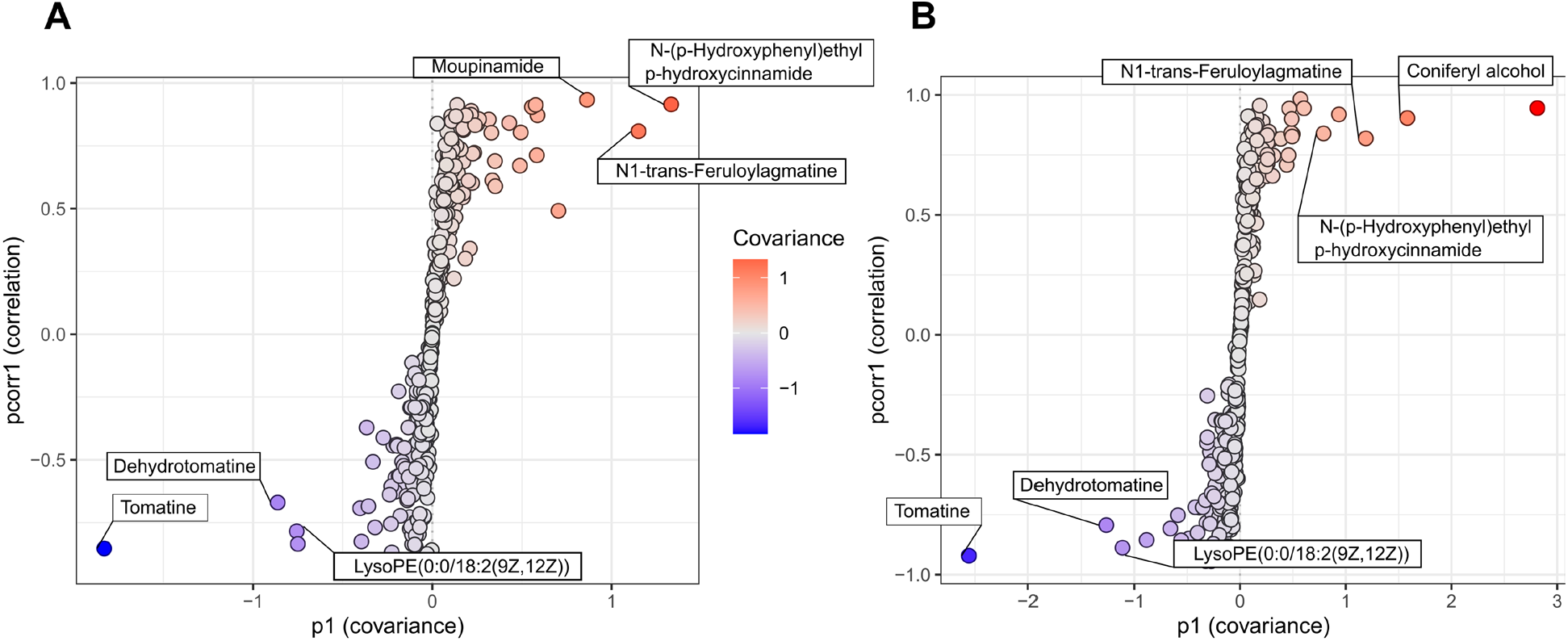
S-plots derived from orthogonal projections to latent structures discriminant analysis (O-PLS-DA) models for the roots of **(A)** AA-treated and **(B)** ANE-treated plants in pairwise comparison with H_2_O across all tested timepoints. Color scale represents covariance strength and direction with red data points indicating most positive covariance and blue indicating most negative covariance.

Bar charts depicting mean LC-MS signals for top OPLS-DA S-plot metabolites visualized across all timepoints illustrate that AA and ANE have similar effects on plant metabolic response **(Fig. 5)**. Treatment of tomato roots with AA and ANE resulted in a sharp increase in metabolic intermediates in ligno-suberin biosynthesis. This includes AA-induced accumulation of moupinamide (syn. N-feruloyltyramine) and significant increases in coniferyl alcohol in the roots of ANE-treated plants across all tested timepoints. In roots, AA and ANE treatments also induced increased levels of N-(p-hydroxyphenyl)ethyl p-hydroxycinnamide and N1-trans-feruloylagmatine compared to H_2_O treatment, reflecting strong upregulation of the shikimate pathway and phenolic compound synthesis. Reduced levels of tomatine and dehydrotomatine were observed in the roots of AA- and ANE-treated plants indicating suppression of steroid glycoalkaloid biosynthesis. Treated plants also showed lower levels of lyso-phosphatidyl ethanolamine (0:0/18:2(9Z,12Z)) that could reflect enhanced membrane lipid turnover.

**Figure 5.**
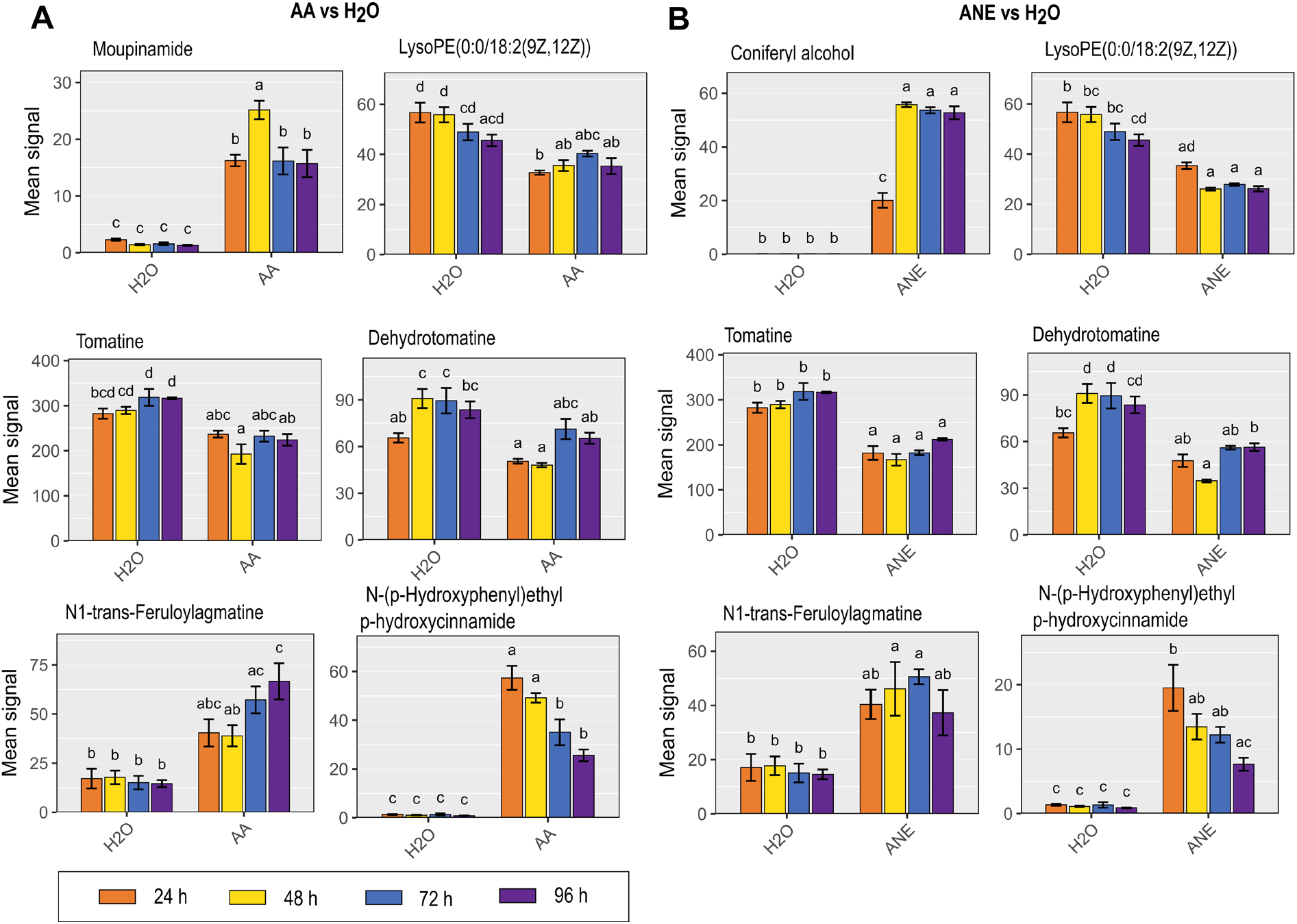
Mean LC-MS signals for the top O-PLS-DA S-plot metabolites visualized across 24, 48, 72, and 96 hour timepoints for **(A)** AA- and **(B)** ANE-treated roots in pairwise comparison with H_2_O.

## Discussion

AA and ANE can induce disease resistance locally and systemically, alter the accumulation of key phytohormones, and change the transcriptional profile of tomato with a striking level of overlap between the two treatments (Lewis et al. 2022). The current study examined and characterized the AA- and ANE-induced metabolomes of tomato. AA and ANE locally and systemically induce metabolome remodeling toward defense-associated metabolic features.

Early studies investigating transcriptional and metabolic changes in potato revealed selective partitioning and shifting of terpenoid biosynthesis from steroidal glycoalkaloids to sesquiterpenes following treatment with AA or EPA or challenged with *P. infestans* (Choi et al. 1992; Stermer and Bostock 1987; Tjamos and Kuc 1982). Similarly, our work here with AA- and ANE-treated tomato seedlings has shown a marked decrease in the levels of two abundant glycoalkaloids, tomatine and dehydrotomatine **(Fig. 5)**. Our data also show strong enrichment of sesquiterpenes in leaves of AA-treated plants at 96 hours post treatment, although the identity of these sesquiterpenes is unresolved **(Supplementary Fig. S2A)**. This work further supports evidence for differential regulation and sub-functionalization of sterol/glycoalkaloid and sesquiterpene biosynthetic pathways in solanaceous plants in different stress contexts (Choi et al. 1992; Stermer and Bostock 1987).

AA and EPA are strong elicitors that are abundant in structural and storage lipids of oomycete pathogens, but absent from higher plants. Although their initial perception by the plant is likely different from that of canonical MAMPs (e.g., flg22, chitosan, lipopolysaccharide), there is some convergence in downstream defenses induced by these various MAMPs. Work to characterize the effect of canonical MAMP treatment on the metabolomes of various plant species has implicated common metabolic changes that prime for defense. Cells and leaf tissue of *A. thaliana* treated with lipopolysaccharide (LPS) showed enrichment of phenylpropanoid pathway metabolites, including cinnamic acid derivatives and glycosides (Finnegan et al. 2016). In the same study, SA and JA were also positively correlated with LPS treatment, as we also observed in tomato following treatment with AA (Finnegan et al. 2016; Lewis et al. 2022).

Recent work in *A. thaliana* wild-type and receptor mutants treated with two chemotypes of LPS showed increases in hydroxycinnamic acid and derivatives and enrichment of the associated phenylpropanoid pathway (Offor et al. 2022). Work in tobacco similarly found treatment with LPS, chitosan, and flg22 all induced accumulation of hydroxycinnamic acid and derivatives, and that defense responses elicited by these MAMPs were modulated by both SA and JA (Mhlongo et al. 2016). More recent work in the cells of *Sorghum bicolor* treated with LPS showed enrichment of hydroxycinnamic acids and other phenylpropanoids in coordination with accumulation of both SA and JA (Mareya et al. 2020). Treatment of tomato with flg22 and flgII-28 also enriched hydroxycinnamic acids, and tomato treatment with cps22 revealed a metabolic shift toward the phenylpropanoid pathway with hydroxycinnamic acid, conjugates and derivatives as key biomarkers (Zeiss et al. 2021a, b). Similar to traditional MAMPs, AA and the AA/EPA-containing complex mixture, ANE, both induce enrichment of cinnamic acid and derivatives in tomato seedlings **(Fig. 3)**. This supports the hypothesis that MAMPs broadly induce similar metabolic changes to enrich pools of specialized secondary metabolites that contribute to plant immunity.

AA- and ANE-treated roots showed strong enrichment of metabolic features classified as fatty acyl glycosides of mono- and disaccharides **(Fig. 3)**. Fatty acyl glycosides have been studied in several plant families and are most extensively characterized in members of Solanaceae (Asai and Fujimoto 2010; Asai et al. 2010; Dalsgaard et al. 2006; Moghe et al. 2017). Investigations into the function of fatty acyl glycosides in plants suggest they may act to protect against insect herbivory through various mechanisms and provide protection against fungal pathogens (Leckie et al. 2016; Luu et al. 2017; Puterka et al. 2003; Simmons et al. 2004; Weinhold and Baldwin 2011). A recent study isolated and identified fatty acyl glycosides from strawberry capable of inducing immune responses in *A. thaliana*, including ROS burst, callose deposition, increased expression of defense-related genes, and induced resistance to bacterial and fungal challenge (Grellet Bournonville et al. 2020). This same work also demonstrated that the strawberry-derived fatty acyl glycosides induced resistance in soybean and, due to their antimicrobial activity, also protected lemon fruits postharvest from fungal infection (Grellet Bournonville et al. 2020). AA- and ANE-root treatments locally elicit accumulation of the same class of defense associated metabolites that Grellet et al. illustrated to have direct antimicrobial activity and protect against disease (Grellet Bournonville et al. 2020).

Cell wall fortification is an important plant defense often initiated upon pathogen infection. Cell wall lignification is a well-studied mechanism with localized accumulation of phenolic intermediates and lignin at attempted penetration sites (Nicholson and Hammerschmidt 1992; Vance et al. 1980; Zeyen et al. 2002). Lignification reinforces and rigidifies the cell wall to create an impervious barrier to microbial ingress (Nicholson and Hammerschmidt 1992; Vance et al. 1980; Zeyen et al. 2002). In our study, AA treatment of tomato roots induced accumulation of a phenylcoumaran intermediate in lignin biosynthesis, while ANE treatment induced accumulation of coniferyl alcohol, an important monomer unit of lignin. Interestingly, coniferyl alcohol has recently been shown to act in a signaling capacity in a regulatory feedback mechanism to intricately control lignin biosynthesis, an irreversible process that is energetically costly (Guan et al. 2022). The findings of our study coincide with the well-characterized role of lignin and its intermediates in plant defense.

This work characterizes local and systemic metabolic profiles of AA- and ANE-treated tomato with the oomycete-derived MAMP, AA, and the AA-containing biostimulant, ANE. AA and ANE profoundly alter the tomato metabolome toward defense-associated secondary metabolites with notable overlap in enriched metabolite classes compared to H_2_O control. Further investigation is required to elucidate the functional contribution of these metabolic features in AA- and ANE induced resistance and, more broadly, plant immunity. Our study adds to the understanding of MAMP-induced metabolomes with implications for further development of seaweed-derived biostimulants for crop improvement.

## Author contributions

Study planned by all authors. Experiments performed by DCL, metabolite analyses conducted by TVDZ and AR. Paper written by DCL, TVDZ, and RMB, with editing by GLC and AR.

## Notes

**Funding:** Research supported in part by a Gates Millennium Scholars Program fellowship and Jastro-Shields awards to DCL, and an unrestricted gift from Acadian Seaplants, LTD., to RMB.

### Competing Interest Statement

The authors have declared no competing interest.

### Summary of Updates

Abstract, results, and discussion sections updated to reflect reviewers comments. New supplemental figure included to reflect reviewers comments.

