## Supplemental Figures for "The oomycete MAMP, arachidonic acid, and an *Ascophyllum nodosum*-derived plant biostimulant induce defense metabolome remodeling in tomato"

### e-*X*tra

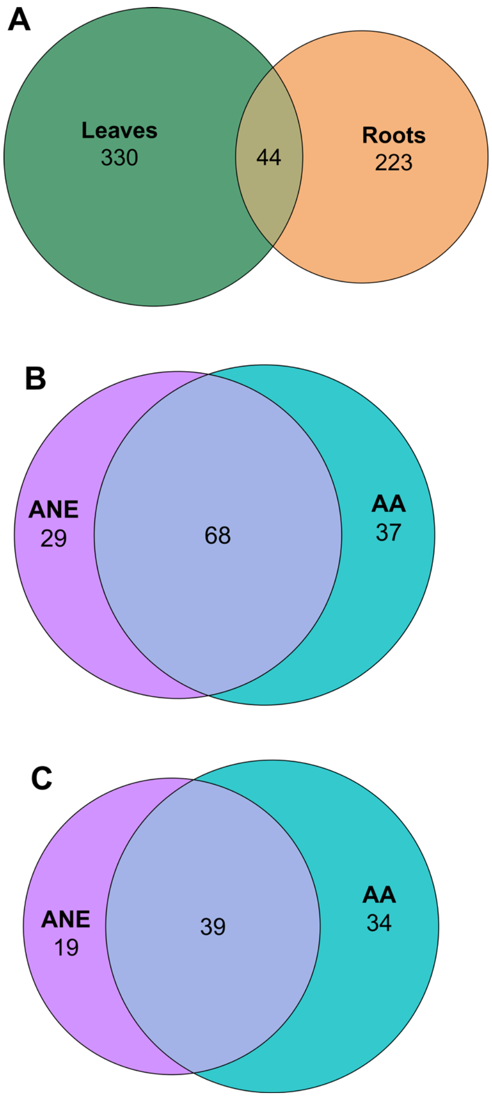


**Supplementary Figure S1**. Venn diagram depicting the comparison of annotated features across all treatments ($H_2O$, AA, and ANE) and timepoints (24, 48, 72, 96 hours) between roots and leaves **(A)**. Venn diagrams depicting the comparison of differential metabolic features of AA- and ANE-treated plants across all tested time points in pairwise comparison to $H_2O$ for roots **(B)** and leaves **(C)** having a fold change >2 and a *p* value < 0.05.

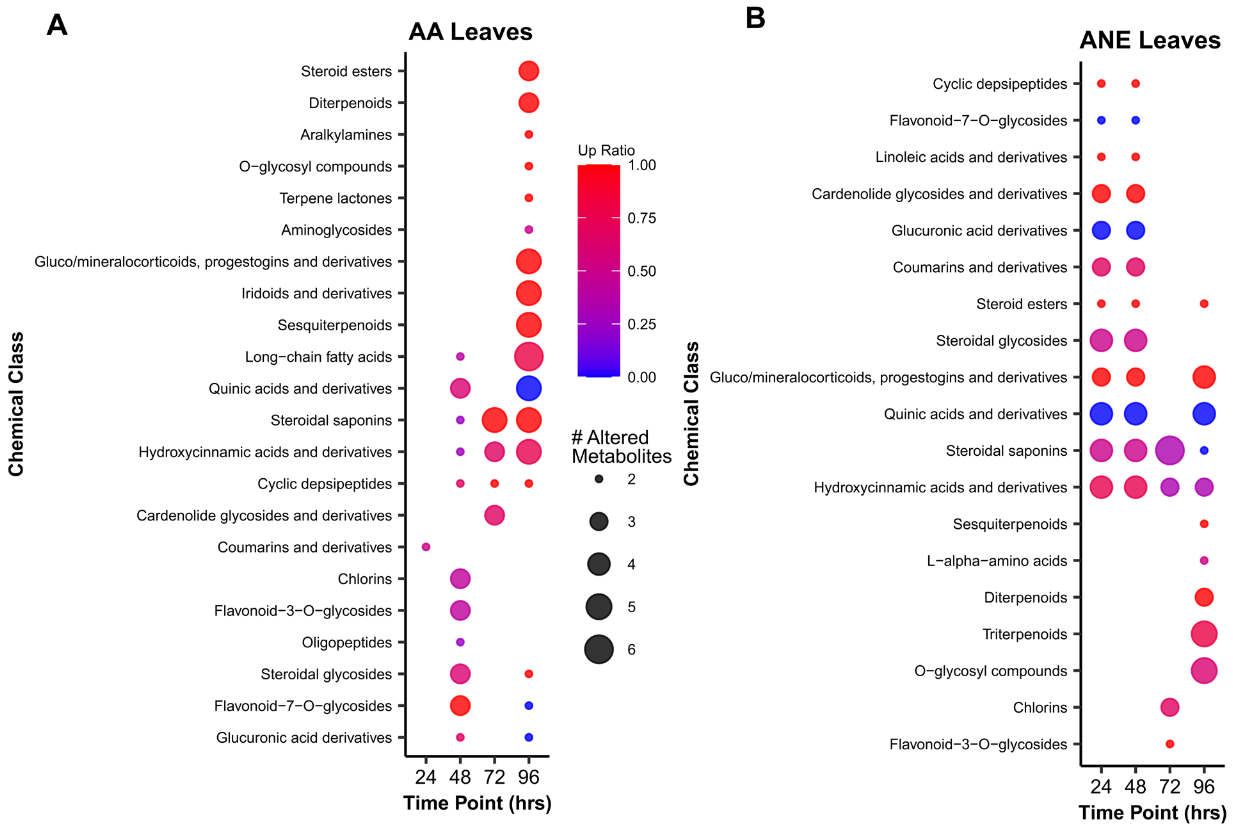


**Supplementary Figure S2**. Chemical enrichment analysis of significantly altered metabolite clusters in leaves of **(A)** AA- and **(B)** ANE-root-treated plants compared to $H_2O$ after 24, 48, 72, 96 hours of treatment. Point sizes indicate the number of altered metabolites per identified cluster. Point color scale indicates the ratio of up (red) and down (blue) compounds in AA- and ANE-treated plants compared to $H_2O$. Up ratio represents the proportion of increased/decreased metabolites compared to $H_2O$.

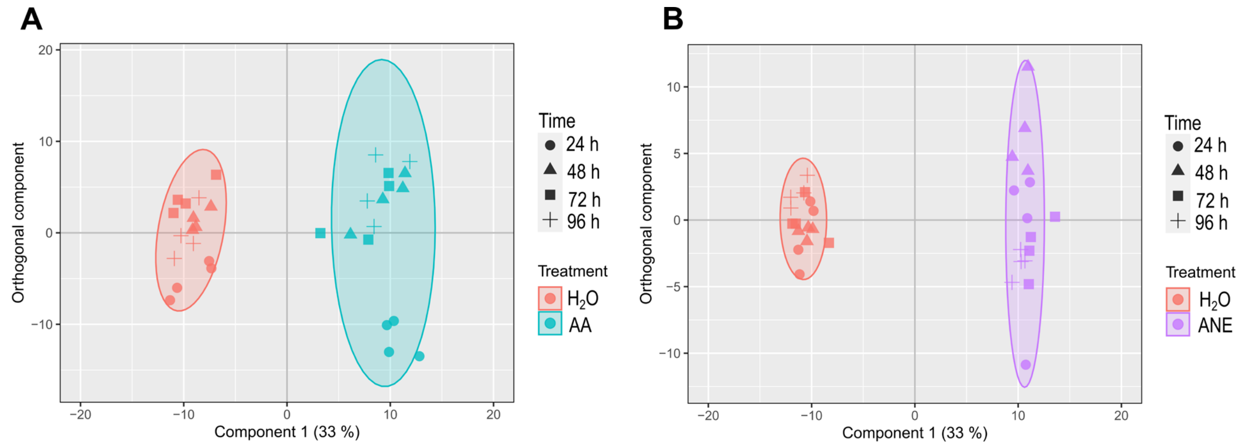


**Supplementary Figure S3**. Orthogonal projections to latent discriminant analysis score plots of tomato root metabolites after 24, 48, 72, and 96 hours of treatment with **(A)** 10 µM AA and **(B)** 0.4% ANE in pairwise comparison to $H_2O$. Ellipses indicate the 95% confidence intervals. Score plots represent four biological replicates performed per timepoint and treatment.
